# Generalized habitual tendencies in alcohol dependent rats

**DOI:** 10.1101/2022.10.04.510642

**Authors:** Francesco Giannone, Arian Hach, Magda Chrószcz, Marion M. Friske, Marcus Meinhardt, Rainer Spanagel, Wolfgang H. Sommer, Anita C. Hansson

**Affiliations:** Institute of Psychopharmacology, Central Institute of Mental Health, Medical Faculty Mannheim, University of Heidelberg, 68159 Mannheim, Germany; Maj Institute of Pharmacology Polish Academy of Sciences, Department of Molecular Neuropharmacology, Krakow, 31-343, Poland

**Keywords:** alcohol addiction, post-dependent animals, goal-directed, habit learning, devaluation, DREADD, natural reward, dorsomedial striatum

## Abstract

Habitual responses and ultimately compulsive behavior are thought to be at the core of addiction including alcohol use disorder (AUD). Little is known whether the habitization concerns exclusively the response towards alcohol or generalizes to other daily activities. Here, we address this question in a well-established animal model of AUD – the postdependent rat model – by testing habitual responses towards a sweet palatable reward in two striatal learning paradigms: spatial navigation and reward conditioning. For the spatial navigation task, alcohol-dependent and control rats were tested on a sequential decision-making test after short and prolonged T-Maze training; for the reward conditioning task, rats were trained under a random interval schedule for a short and prolonged period and tested in a satiety devaluation test at each time point. Another cohort of alcohol-naive rats was trained and tested on both paradigms under DREADD (designer receptors exclusively activated by designer drugs)-mediated inactivation of the dorsomedial striatum (DMS) which controls goal-directed behavior. Our results show that alcohol-dependent rats displayed increased habitual behavior to obtain saccharin reward on both paradigms, with overall more habitual choices after prolonged training on the spatial navigation task, and increased habitual responses already after short training on the reward conditioning task. Finally, DREADD-mediated inactivation of the DMS increased habitual behavior in non-dependent rats on both paradigms. Our results provide evidence that a history of alcohol dependence produces a bias towards habitual responding that generalizes to a natural reward in rats. Similarly, a habitual bias was induced in non-dependent rats after inactivation of the DMS, thus confirming the critical role of this region in maintaining goal-directed behavior and suggesting its diminished control in AUD.

## Introduction

A habit is a consolidated response to a stimulus, which once acquired is relatively resistant to interference and as such largely independent of action-outcome contingencies (Yin and Knowlton, 2006; Yin et al., 2009). A habit develops by extensive stimulus-response pairings and repetition of the same response sequence in the same context, and it is thought to depend on striatal learning processes. There is considerable evidence, mostly coming from rodent studies, suggesting the involvement of two different learning processes, the dorsomedial striatum (DMS) controlling the acquisition of goal-directed actions mainly via dopamine D1 receptors, and the dorsolateral striatum (DLS) acquisition of habits and skills via dopamine D2 receptors (Yin et al., 2009; Sommer et al., 2014). The two striatal regions receive excitatory inputs from different areas of the cortex. The DMS receives modulatory input from prefrontal cortices, whereas the DLS primarily receives major inputs from sensorimotor and premotor cortices in both humans and rats (Balleine et al., 2009; Balleine and O’Doherty, 2010; Corbit and Janak, 2016a). Dependent on DMS activity, goal-directed behavior is mainly involved during the early instrumental conditioning, and with overtraining habitual responding occur under control of the DLS (Yin et al., 2004, 2005a; Yin et al., 2005b; Yin and Knowlton, 2006; Yin et al., 2006; Balleine et al., 2009). Some studies suggest that the DMS and DLS operate in parallel during the transition from goal-directed to habitual responses (Yin et al., 2009; Vandaele et al., 2019), with the DLS already active at the early onset of training (Kupferschmidt et al., 2017; Bergstrom et al., 2018; Smith et al., 2021), and it appears that both subregions act via an opposing and competitive relationship (Smith and Graybiel, 2013; Turner et al., 2022).

The harmful use of alcohol is a global problem causing 3 million deaths per year (which accounts to 5.3% of deaths worldwide) and is attributed to the world’s third largest risk factor for premature mortality, disability and loss of health (Rehm et al., 2009; WHO, 2018). When considering the total harm inflicted by alcohol including its societal costs, alcohol is by a wide margin the most harmful drug in the western world (Nutt et al., 2010). Alcohol use disorder (AUD) is characterized by the loss of flexible control over drug use despite negative consequences (Rehm et al., 2009; Connor et al., 2016; Koob, 2021). Current addiction theories postulate that the loss of control may be explained with increased habitual behavior in which drug use persists even when the drug is no longer rewarding, and ultimately leading to compulsive alcohol drinking (Everitt and Robbins, 2005; Belin et al., 2009; Belin et al., 2013; Everitt and Robbins, 2016) and habitual drug-seeking (Everitt et al., 2008; Ostlund and Balleine, 2008; Belin et al., 2009; Root et al., 2009). Recent research has also implicated the dorsal striatum in the control of decision-making regulated by reward, particularly the role of the DMS in the integration of reward-related processes with action control (Balleine et al., 2007). The DLS has been linked with habitual responding for alcohol in animals (Corbit et al., 2012; Serlin and Torregrossa, 2015; Giuliano et al., 2019) and natural rewards (Dickinson et al., 2002; Corbit et al., 2012; Houck and Grahame, 2018; Turner et al., 2022) and humans (Vollstadt-Klein et al., 2010).

Most studies assessing habitual responses in animals use operant conditioning, and there are only few publications demonstrating the impact of alcohol exposure. Both acute (Houck and Grahame, 2018) and chronic ethanol exposure (Corbit et al., 2012; Renteria et al., 2018; Towner and Spear, 2021) shifted animals toward behaving more habitually. Most of these studies assess habitual responses based on their sensitivity to devalued reinforcers. The results are difficult to interpret, since they rather may relate from impaired instrumental learning abilities than habits (Balleine and Dezfouli, 2019).

In rodents, the spatial navigation task has been successfully used for assessment of goal-directed and habitual behavior (Packard, 1999; Yin and Knowlton, 2004; Lee et al., 2008; McCracken and Grace, 2013). Thus, spatial navigation could present an alternative to reward conditioning, although there are currently no data available from spatial navigation tasks assessing alcohol-related habitual responses in rodents.

This study aimed to investigate how habit formation processes are affected by a history of alcohol dependence using two striatal learning paradigms, the spatial navigation and reward conditioning task. We tested habitual responses towards a sweet palatable reward in both tasks in a well-established animal model of AUD – the post-dependent (PD) rat model. Alcohol dependence was introduced in adult rats by repeated cycles of intermittent exposure (CIE) to alcohol vapor intoxication and withdrawal (Rimondini et al., 2002). We further analyzed the effects of a designer receptors exclusively activated by the designer drugs (DREADD)-mediated inactivation of the DMS in habitual behavior.

## Materials and Methods

### Experimental animals and design

Male Wistar rats were obtained from Envigo (Germany), 6 weeks of age at arrival. Rats were group-housed (four animals per Makrolon type 4 cage, cage dimension: 85 × 85 × 67 cm) under a 12h light/dark cycle (lights off at 6 AM, lights on at 18:00 PM) with ad libitum access to water and food, and left to acclimatize to the environment for one week. They were handled daily for the following week before beginning the experimental procedures. All experimental procedures started in the dark phase (2hrs after lights off) in adult rats (≥ 8 weeks). The experimental procedures were approved by the Regierungspräsidium Karlsruhe, Germany (license: G-166/21).

Three separate cohorts of male rats were used for this research, each allocated to a different experiment. Study 1 was performed to assess the effects of alcohol dependence on habitual behavior in the context of spatial navigation (Experiment 1) and reward learning (Experiment 2) tasks (**Figure 1**). Study 2 was performed to evaluate the effects of DREADD-mediated posterior dorsomedial striatum (pDMS) inactivation on habitual behavior in the context of both spatial navigation and reward learning tasks (Experiment 3, **Figure 1**).

**Figure 1.**
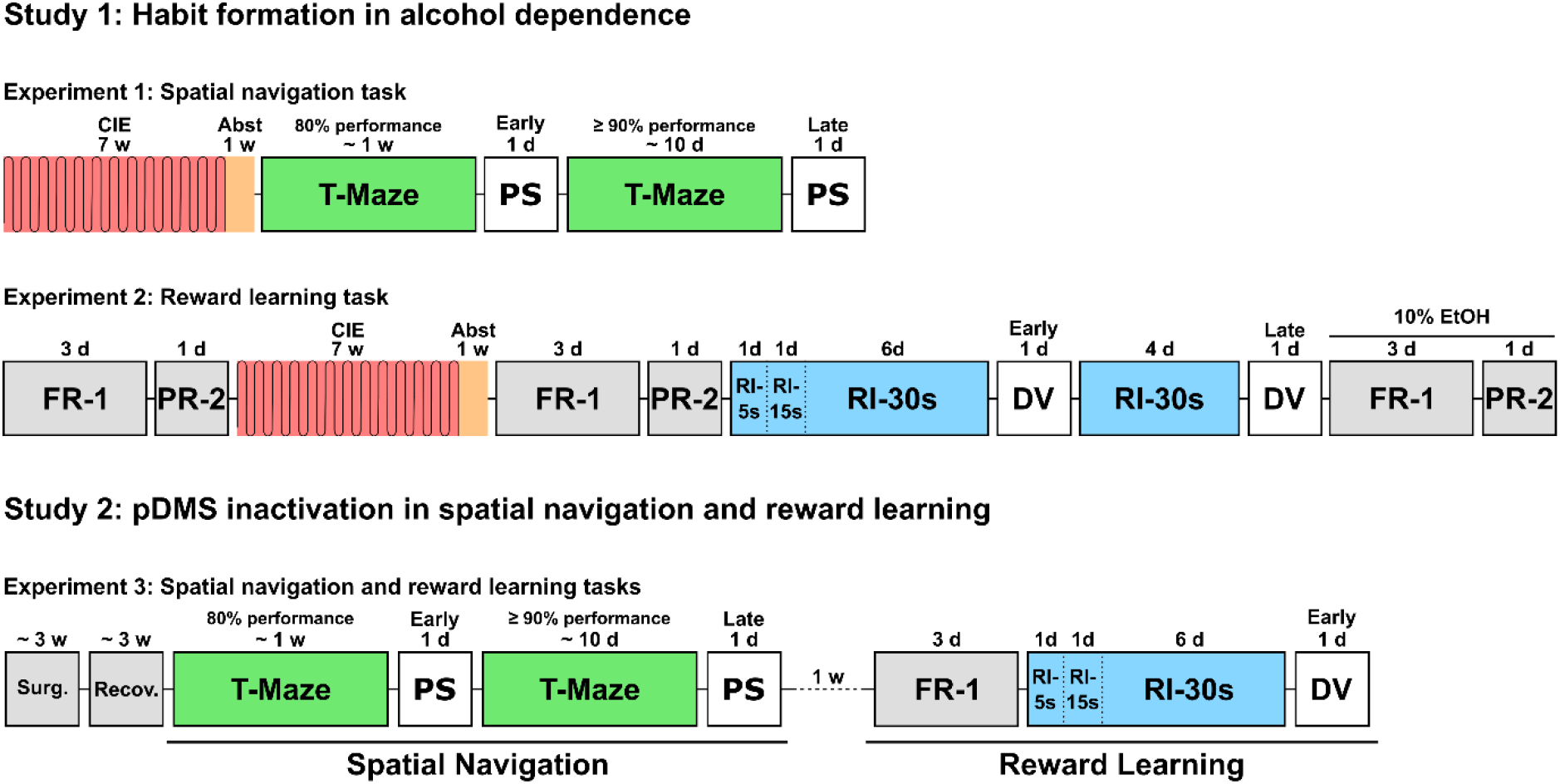
Experimental timelines. In Study 1 habitual responses (spatial navigation, reward learning) were analyzed in two cohorts of alcohol dependent and non-dependent rats. At the end of experiment 2 rats were trained to self-administer 10% ethanol to confirm the alcohol dependent phenotype. In Study 2 rats were tested for both behavior paradigms after inactivation of the posterior part of the dorsomedial striatum (**pDMS**). In all three experiments 0.2% saccharin was given as a natural reward. **CIE:** Chronic Intermittent Ethanol vapor exposure, **Abst:** abstinence, **FR-1:** Fixed Ratio 1 operant schedule, **PR-2:** Progressive Ratio-2 operant schedule, **RI:** Random Interval operant schedule (5s, 15s, and 30s indicate the interval length in seconds), **PS:** probe session composed of 5 consecutive trials, **DV:** Devaluation test, **Surg.**: surgery, **Recov.**: recovery from surgery.

### Material details

#### Chemicals, drugs and kits

Alexa Fluor 488 donkey anti-mouse IgG(H+L) (Thermo Fisher, Ref: AB_141607), Alexa Fluor 594 donkey anti-rabbit IgG(H+L) (Thermo Fisher, Ref: AB_141637), anti-fade mounting medium (Shandon Immu-Mount^™^, Thermo Scientific), Carprofen (50 mg/ml, Zoetis Deutschland GmbH, Germany), Clozapine N-oxide (CNO) dihydrochloride (BIOZOL Diagnostica Vertrieb GmbH, Germany), DREADDs (ssAAV-8/2 virus expressing either hM4D(Gi)-mCherry (titer: 7.3 × 10^12^) or eGFP (titer: 6.9 × 10^12^), University of Zürich and ETH Zürich, Neuroscience Center Zürich ZNZ, Viral Vector Facility VVF, Switzerland), isoflurane (1 ml/ml, CP-Pharma Handelsgesellschaft GmbH, Germany), ethanol (96% vol, VWR BDH Chemicals, France), ketamine (bela-pharm GmbH & Co. KG, Germany), xylazine (Wirtschaftsgenossenschaft deutscher Tierärzte eG, Germany), kit for measurements of blood alcohol concentration (BAC, containing Alcohol buffer pH 7.4, alcohol oxidase (AOD) and AM1 analyzer, Analox Instruments Ltd, United Kingdom), lidocaine (2%, beta-pharm GmbH & Co KG, Germany), mouse anti-GFP (Invitrogen, Ref: A22230), paraformaldehyde (PFA, Carl Roth GmbH & Co. kg, Germany), rabbit anti-RFP (Rockland, Ref: 600-401-379), saccharin (saccharin sodium salt hydrate, Sigma-Aldrich Chemie GmbH, Germany), Sodium azide (Merk KGaA, Germany).

### Method details

#### Induction of alcohol dependence

Alcohol dependence (Experiments 1 and 2, **Figure 1**) was induced via chronic intermittent ethanol (CIE) vapor exposure (Rimondini et al., 2002; Hirth et al., 2016; Meinhardt et al., 2013). The vapor exposure was conducted into custom-made vapor chambers that could hold four Type-4 cages each. After one week of habituation to the chambers, the vapor exposure procedure was initiated. The rats were exposed to alcohol vapors for 16 hrs per day every day for 7 weeks. The exposure was initiated 4 hrs before the beginning of the active cycle and ended at the end of the active cycle. The ethanol was delivered by dosing pumps into stainless-steel coils (Knauer, Berlin, Germany) that were heated to 70°C and released into the chambers at a rate of 16 L/min, mixed with air. Twice per week, right after the end of the vapor exposure session, 4 animals per chamber were randomly picked to measure BACs. The pump rate was adjusted in order to keep the BACs between 150 and 300 mg/dL throughout the procedure. At the end of the seventh week of exposure, the animals were maintained into abstinence (air exposure only) for one week before initiating the behavioral trainings. Seven hours after the last ethanol vapor exposure cycle each rat was monitored for up to 5 min to assess tremor, tail rigidity, vocalization, abnormal gait, and ventral limb retraction. Severity of each withdrawal sign was ranked as 0 (non-existent), 1 (mild), and 2 (severe) added up and expressed as total withdrawal scores (Macey et al., 1996; Uhrig et al., 2017).

#### Spatial navigation (T-Maze) task

The spatial navigation task was performed in a custom-made T-Maze. The maze was elevated 50 cm above the floor and had four closed identical arms (50 cm (L) × 10 cm (W) × 50 cm (H). The access to either the north or the south arm was blocked by using a 9.8 cm cuboid. The training and testing protocols for habitual behavior were adapted from Packard & McGaugh (1996) and Yin & Knowlton (2004). On the first day, the animals underwent a habituation session. The access to the north arm of the maze was blocked; each rat was positioned at the end of the south arm and was free to explore the maze for 5 minutes. On the following day, the training started. The access to the north arm was still blocked, on the wall at the end of the west (left) arm a visual cue was positioned (baited arm) while no optic cue was presented at the east (right) arm (unbaited arm). At the end of both side arms a transparent plastic well was attached on the floor of the maze. The well at the west arm always contained 1 ml of 0.2% saccharin solution, while the well at the east arm was always kept empty.

For both Experiment 1 and 3, each training trial started by positioning the rat at the end of the south arm and ended after 5 minutes or after the animal found and consumed the reward, after which the animal was removed from the maze. Each training session consisted of 5 consecutive trials. For each trial, the performance of the animal was scored: A correct choice was considered when the animal walked from the south arm directly into the baited arm and consumed the reward, in all other cases (the animal walked into the baited arm but did not consume the reward or it walked into the unbaited arm) the choice was considered as incorrect. Each animal was trained daily (one session of 5 consecutive trials per day) until 80% performance (four correct choices out of 5 trials) for two consecutive sessions was reached, to ensure the animals properly learned the task. Then a probe session was performed on the following day (Early time point, Figure 1). After the early probe session, the training resumed and was continued daily until a criterion of at least 90% performance over the course of 10 consecutive days was reached, to ensure overtraining, after which a second probe session was performed (Late time point, Figure 1).

The probe sessions consisted of 5 consecutive probe trials, each trial was identical to the training trials except that animals were inserted into the north arm of the maze while the south arm was blocked; the animal was removed from the maze immediately after making a choice (baited or unbaited arm) and walked back into the center of the maze. When the animal walked into the baited arm, this was considered as a goal-directed choice. When the animal walked into the unbaited arm it was considered as a habitual choice.

#### Reward learning (operant conditioning) task

Before initiating the training sessions, all animals from experiments 2 and 3 underwent a 30 minutes habituation session in which no lever and no cue was present, the reward (0.2% saccharin, 13 μl drop) was released into the receptacle at a random interval 60s (RI-60s) schedule.

Following the habituation, the animals from Experiment 2 (**Figure 1**) were trained under fixed ratio 1 (FR-1) schedule until they reached a criterion of at least 50 rewards for 3 consecutive sessions, after which, a progressive ratio 2 (PR-2) test was performed. Based on the breakpoint value, the animals were then pseudo-randomly assigned to the treatment groups (CIE: chronic intermittent ethanol vapor exposed group, air-exposed control group) and the vapor exposure was initiated. One week after the last ethanol vapor exposure, the animals underwent three more sessions under FR1 schedule and were again tested in a PR-2 test. After the last PR test, all animals underwent one session under random interval 5s (RI-15s) schedule, one session under RI-15s schedule, and 6 sessions under random interval 30s (RI-30s) schedule. The training was performed from Mondays to Thursdays. After the 6^th^ day of RI-30s training the animals underwent the devaluation (DV) test with a satiety devaluation procedure of 30 min followed by a 5 minutes extinction test (Early time point). The following week the RI-30 training was resumed for another 4 sessions and the DV test was repeated (Late time point).

For Experiment 3 (**Figure 1**) the animals were also pre-trained for three days under FR-1 schedule, after which they were directly trained under random interval schedule (RI-5s, RI-15s, RI-30s) and tested in a devaluation test at the Early time point only.

FR-1 pre-training sessions in Experiment 2 ended after 30 minutes, FR-1 pre-training sessions in Experiment 3 ended after 30 minutes or after 50 rewards were earned.

All random interval sessions ended after 30 minutes or after 30 rewards were earned. Animals that did not earn 30 rewards in the last two sessions prior to the DV test were not included for statistical analysis.

During the DV test all animals from Experiment 2 and 3 were put into single cages and brought into a novel environment, where they were given one bottle containing 0.2% saccharin and one bottle containing tap water, and they were allowed to freely drink for 30 minutes. Immediately after the end of the drinking session, all animals were brought into the operant boxes where they underwent a 5 minutes extinction session in which the house light was turned on, the lever was present but no reward was released. At the end of the extinction session all animals were brought back into the single cages and were given another 30 minutes of free drinking to confirm the satiated state.

In Experiment 3, 30 minutes prior to each probe session (spatial navigation) and to the extinction session (reward learning), all animals received a 10 mg/ml CNO ip injection (in saline).

#### Progressive ratio test

Prior to the beginning and one week in abstinence all animals in experiment 2 were tested in a PR-2 test. This test provides an estimate about the motivation for an animal to acquire the reward. During the test only the active lever was available, this was the same lever used for the FR1 and the RI trainings. In a PR schedule the number of lever presses (n) required for obtaining the reward increased with each reinforcement (1, 2, 4, 6, 8, 10, … n+2) within a session. The session ended after 2 hours or after no lever press was performed for three consecutive minutes. During the test the house light signaled the beginning and the end of the session. The maximum number of lever presses for a single reinforcement within a session was considered as breakpoint.

#### Alcohol operant training

The animals that completed the reward learning task in experiment 2 were further trained to self-administer 10% ethanol in tap water. The training sessions were identical to the previous saccharin training sessions except that the lever assigned to the ethanol was the opposite lever in the operant chamber. The reward was delivered on the receptacle next to the active lever, on the opposite side compared to the receptacle and the lever used for saccharin training. The animals were trained until they earned at least 50 ^ rewards for three consecutive sessions, after which they were tested in a PR2 test that ended after 2 hours or after 3 minutes of inactivity (see **Figure 1**).

#### Stereotaxic surgeries

Serotype 8/2 adeno associated viral vectors (ssAAV-8/2) expressing either hM4D(Gi)-mCherry (titer: 7.3 × 10^12^) or eGFP (titer: 6.9 × 10^12^) under the control of human synapsin promoter (hSyn1) were used. Surgeries were performed as described in Broccoli et al. (2018) and Meinhardt et al. (2022). Inactivating DREADD virus was injected bilaterally into the pDMS (coordinates: A/P = −0.4; M/L = ± 2; D/V = −4) using 10 μl NanoFil syringes equipped with 33 gauge beveled NanoFil needles (World Precision Instruments, Inc.) at a volume of 0.7 μl and a rate of 0.1 μl per minute. Before activating the micropump (MicroSyringe Pump Controller, Micro 4 ^™^, World Precision Instruments, Inc.) for the injection, the needle was first brought into position based on the aimed coordinates and left in place for 5 minutes. After injection, the needle was kept in place for another 10 minutes before slowly withdrawing it.

Four% isoflurane-oxygen mixture was used for induction of anesthesia, 2-2.5% isoflurane-oxygen mixture was used for maintenance (flow rate: 1 L/min). Carprofen (5 mg/kg, sc) and Lidocaine (1% sc, under the scalp) were injected at beginning of the surgery, Carprofen treatment was continued for three days post-surgery.

#### Perfusions

At the end of the experimental procedures, all rats that received bilateral ssAAV-8/2 injection were perfused. Briefly, rats were deeply anesthetized with a mixture of ketamine-xylazine (100 mg ketamine/kg, 3 mg xylazine /kg, ip), they were then perfused with 150 mL of 1xPBS followed by 100 ml of ice-cold 4% PFA (flow rate: 15 mL/min). The brains were then removed and post-fixed into 4% PFA at 4°C for 48 hours before being transferred into 1 × PBS + 0.1% sodium azide (4°C).

#### Immunofluorescence

For immunofluorescence, consecutive coronal sections (50 μl thickness) were obtained using a vibrating-blade microtome (Leica VT1000 S, Germany) covering the entire dorsal striatum. The slices were then permeabilized into 1% Triton X-100 in 1 × PBS (30 minutes, 4°C), washed three times into 1 × PBS (5 min each, RT), blocked with 10% donkey serum in 1 × PBS (1 hour, RT), and incubated o/n with either 1:1000 mouse anti-GFP (Invitrogen, Ref: A22230) or 1:1000 rabbit anti-RFP (Rockland, Ref: 600-401-379). The slices were then washed three times into 1 × PBS (5 min each, RT), incubated with either 1:500 Alexa Fluor 488 donkey anti-mouse or 1:500 Alexa Fluor 594 donkey anti-rabbit for 1 hour, RT. They were finally washed three times in 1 × PBS (10 min each, RT) before being mounted on glass slides with anti-fade mounting medium. The expression of both eGFP and mCherry was detected by an epifluorescent microscope (Zeiss, Axio Imager M2, Germany).

#### Data analyses and statistics

For the spatial navigation task, two different variables were used to assess habitual behavior: the number of habitual choices within the first trial of each probe session and the overall choice performance calculated as performance ratio: number of goal-directed choices per session divided by the total number of trials per session.

For measuring habitual behavior in the reward learning task, the cumulative devaluation index was used. This was calculated as follows:

(baselinecumulative — testcumulative) / (baselinecumulative + testcumulative)

“Baseline_cumulative_” represents the cumulative average number of lever presses performed for each 1-minute bin within the first 5 minutes of the last two sessions prior to the test; “test_cumulative_” represents the number of lever presses performed for each 1-minute bin within the first 5 minutes of the test. Using the cumulative devaluation index it is possible to analyze how the behavioral approach of the animals (goal-directed or habitual) changes over the course of the test.

The difference between groups in the spatial navigation task using the number of habitual choices within the first trial of each session was analyzed via χ^2^ test. Habitual behavior in the reward learning task was assessed via one-sample t-test of the devaluation index against 0.

The withdrawal scores were compared between groups via Mann-Whitney U test. The remaining data were analyzed via Repeated Measure Mixed Model (Gueorguieva and Krystal, 2004). The estimation method used was Maximum Likelihood (ML), the covariance matrix was chosen based on the lowest BIC. When applicable, LSD post-hoc test was performed. For statistical analysis IBM SPSS Statistics 27 was used.

#### Sample size

Experiment 1: 8 Ctrl and 8 CIE animals were used for the spatial navigation task. One animal died during the vapor exposure, therefore the final sample size was 8 Ctrl and 7 CIE.

Experiment 2: For the reward learning task, 26 rats were used in total (12 Ctrl + 14 CIE). Of these, 2 rats died during the vapor exposure, 3 controls and 3 CIE were excluded from the analysis because they did not reach the 30 rewards per session criterion during the random interval training. The final sample size was 12 Ctrl and 9 CIE.

Experiment 3: A total of 28 rats were used, 14 GFP and 14 hM4Di injected rats. Of these, 2 hM4Di rats were excluded due to misplacement of injection site, 3 hM4Di rats did not acquire the lever pressing behavior, 2 Ctrl and 3 hM4Di were excluded from the analysis, because they did not reach the 30 rewards per session criterion during the random interval training. The final sample size was 14 GFP and 12 hM4Di for the spatial navigation task, and 12 Ctrl and 7 hM4Di for the reward learning task.

## Results

### Study 1: Habit formation in alcohol dependence

Habitual and goal-directed behavior was determined in alcohol dependent and nondependent rats using spatial navigation (T-Maze, Study 1, Experiment 1) and reward conditioning (operant boxes, Study 1, Experiment 2) tasks with 0.2% saccharin as reward. Dependence was introduced by chronic intermittent cycles of ethanol vapor exposure (CIE, (Rimondini et al., 2002; Hirth et al., 2016). Average BAC per daily cycle and experiment was 269.72 ± 16.04 mg/dL. Dependence was further confirmed by measurement of withdrawal scores 7 hrs in abstinence after the last vapor exposure cycle. The average total withdrawal score was 4.38 ± 0.36 for CIE animals compared to control rats (1.65 ± 0.22, U = 22.50, z = - 4.60, p < .000).

### Experiment 1: Spatial Navigation (T-Maze) task

The training on a T-Maze started in one week abstinent rats. Rats had to navigate to a well containing the reward (0.2% saccharin solution) signaled by a visual cue, always located on the left arm of the maze. Each rat was tested individually on different days depending on its performance. For the Early time point each rat was tested in a probe session the following day after making at least 4 correct choices out of five trials (80% performance) for two consecutive sessions. For the Late time point rats were tested after making 10 sessions with an overall performance of at least 90%.

As expected, we found no difference in terms of training performance between groups (F_1,25_ = .02, p = .878) but the performance significantly differed between time points (F_2,48_ = 117.5, p = .000; **Figure 2A**). The post-hoc test indicated that prior to the early probe session the animals properly learned the task as their performance increased compared to the beginning of the training (Start vs Early: p= .000), and prior to the Late probe session the animals improved their performance even more (Early vs Late: p=.001).

**Figure 2.**
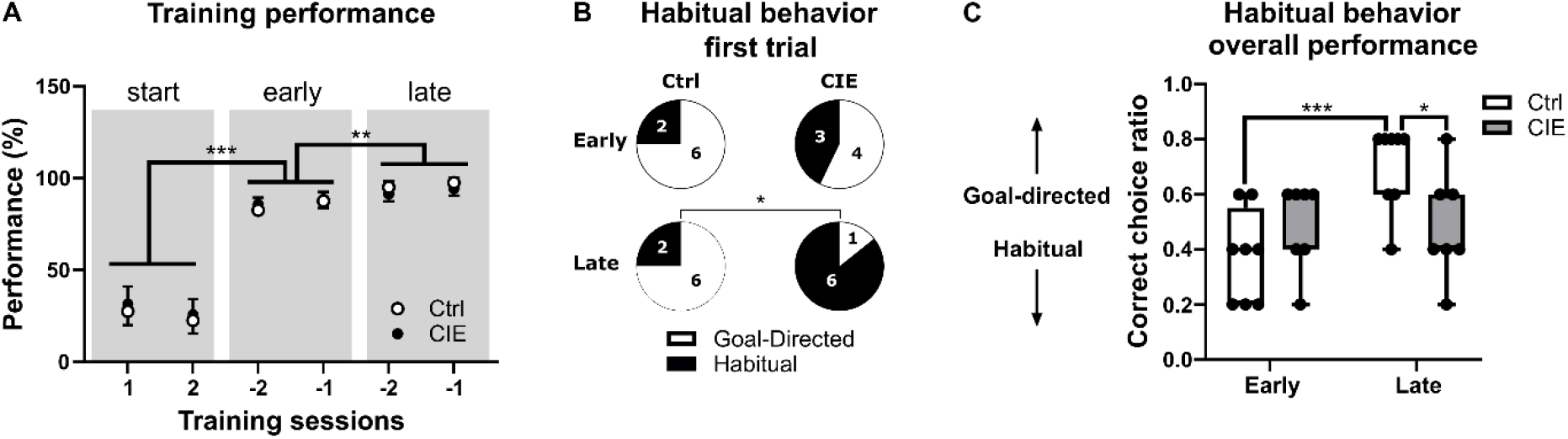
Pronounced habitual behavior in alcohol dependent rats (Study 1, Experiment 1). **A.** Training performance in percentage during the first two sessions, the last two sessions before the Early probe session, and the last two sessions before the Late probe session. Data are shown as mean ± SEM **B.** Pie chart showing number of habitual choices during the first trial of each probe session. **C.** Overall performance including all five trials of Early and Late probe session. Data for correct choice ratio are shown as box and whisker plot (median, first and third quartile range). Data in **B** were analyzed by χ^2^ test. Data in **A** and **C** were analyzed by Repeated Measure Mixed Model, *p < 0.05, **p < 0.01, ***p < 0.001. CIE, cyclic intermittent alcohol vapor exposed rats; Ctrl, control rats. For Details, see Material & Methods.

The results of the probe sessions, considering the first trial of each session (**Figure 2B**) and the overall performance for each session (**Figure 2C**) indicate that CIE animals displayed an increased habitual behavior response compared to controls. Based on the first trials, CIE animals made significantly more habitual turns compared to controls at the Late time point (*χ*^2^ = 5.5, p = .019), 86% CIE vs 25% Controls. From the Early to the Late time point the number of CIE animals that made habitual turns increased (from 43% to 86%) but such increase was not significant (χ^2^= 2.8, p = .094). However, control animals did not show any change in their choices with overtraining as at both time points only two animals made a habitual turn (**Figure 2B**). Based on the overall performance, measured as correct choice ratio per time point, the CIE animals showed a reduced performance, hence an overall increased number of habitual turns, specifically at the Late time point after overtraining (LSD: p = .013; Group*Time interaction: F_1,30_ = 8.1, p = .008). Control animals showed an increased overall performance from the Early to the Late probe session (p=.000) which was not the case for the CIE group (p=1).

### Experiment 2: Reward learning (operant conditioning) task

In order to study habitual behavior in the context of reward learning, we first trained male Wistar rats to self-administer 0.2% saccharin under a FR-1 schedule until they obtained a minimum of 50 rewards for three consecutive sessions after which their motivation for the reward was tested in a progressive ratio PR-2 test. When analyzing the number of lever presses during the FR1 sessions prior to each PR-2 test, we found a significant Group × Time × Session interaction (F_2,168_ = 5.1, p=.007). The post-hoc test showed no difference before initiating the vapor exposure (**Figure 3A**).

**Figure 3.**
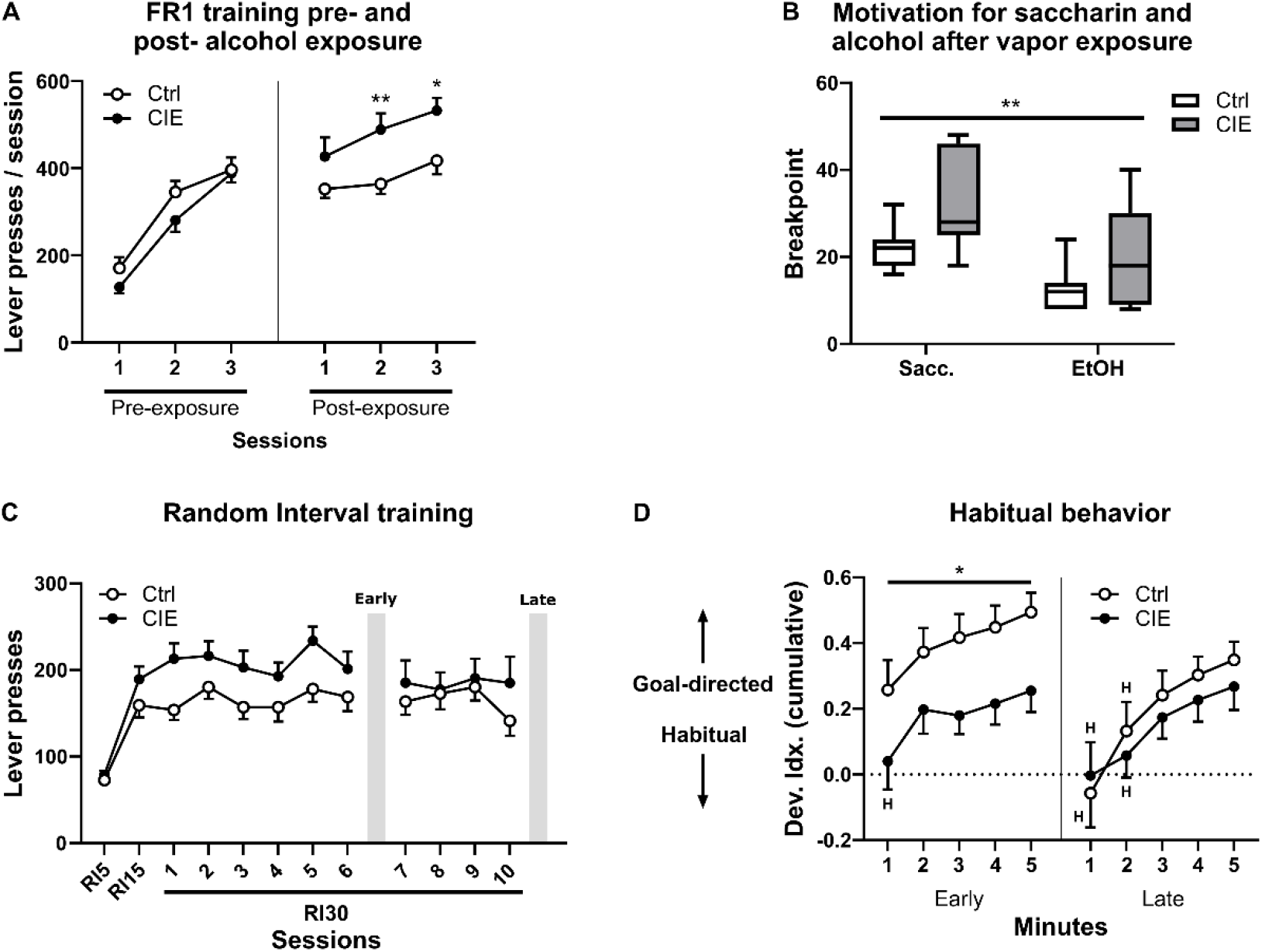
Alcohol dependent rats displayed habitual behavior for reward conditioning (Study 1, Experiment 2). **A.** Lever presses during FR1 training before and one week after the last cycle of ethanol vapor exposure. **B.** Breakpoint for 0.2% saccharin and 10% ethanol. **C.** Lever presses during random interval training. **D.** Devaluation index (Dev Idx) over the course of the extinction test for both early and late time-points. A dev Idx of 0 indicates full habitual behavior; a Dev Idx of 1 indicates full goal-directed behavior. Data points highlighted with *“H”* are indicating habitual behavior. Data in graphs **A**, **C** and **D** are shown as mean ± SEM. Data in graph **B** are shown as box and whisker plot (median, first and third quartile range). Data were analyzed via Repeated Measure Mixed Model. *p < 0.05, **p < 0.01, ***p < 0.001. CIE, cyclic intermittent alcohol vapor exposed rats; Ctrl, control rats. For Details, see Material & Methods.

We used the breakpoint (BP) value obtained from this test to pseudo-randomly assign the animals into the two treatment groups (CIE and Ctrl) in order to control for the motivational level toward the reward. The mean BP for the two groups was 25.1 ± 2.1 and 26.2 ± 2.0 (mean ± SEM) for the prospective Ctrl and the CIE groups, respectively (t_19_ = −.3, p = .731).

All animals were then tested again for 0.2% saccharin (on the same lever and receptacle) 1 week after the last cycle of ethanol vapor exposure. The CIE group showed significantly increased responding for saccharin in the last two sessions (2^nd^ session: p=.002, 3^rd^ session: p=.028). To confirm the alcohol dependent phenotype, at the end of the whole experiment animals were trained to self-administer 10% ethanol (on the opposite lever and receptacle). The BP obtained from the PR-2 tests for 0.2% saccharin and 10% alcohol, indicated that CIE animals displayed a higher motivation for both rewards compared to controls (F_1,40_ =13.3, p = .001), but both groups strongly preferred saccharin over alcohol (effect of reward: F_1,40_ = 16.0, p=.000) (**Figure 3B**).

After the second PR-2 test for saccharin, habitual behavior was induced by training the rats for one session under RI-5 schedule, one session under RI-15 schedule and for 10 sessions under RI-30 schedule. Despite the higher motivation for saccharin observed in the CIE animals, the two groups did not differ significantly in terms of lever presses during the Random Interval (RI) training (F_9,93_ = 3.8, p = .081) (**Figure 3C**).

Importantly, also the baselines used for calculating the devaluation index did not differ between groups, as indicated by the lack of a Group effect (F_1,28_ = .1, p = .751) or any interaction involving this factor.

Habitual and goal-directed behavior was tested after six RI-30s sessions (Early time point) and after four additional RI-30s sessions (Late time point). The test session (5 min) was performed under extinction condition and commenced immediately after 30 minutes of satiety devaluation. When analyzing the cumulative devaluation index of the two groups, we found a significant Group × Timepoint interaction (F_1,135_ = 5.8, p = .018). The post-hoc test revealed that at the Early time point CIE animals had significantly lower devaluation index compared to controls (p = .011) while the groups did no longer show any difference at the Late time point (p = .432) (**Figure 3D**). Moreover, control animals significantly reduced their devaluation index from Early to Late time point (p = .000), while CIE animals did not change between the two time-points (p = .494) (**Figure 3D**). Finally, we found a significant main effect of Bin (F_4,52_ = 10.1, p = .000) with no interaction involving this factor, indicating that both groups, at both time points progressively became aware of the lack of reward during the test, correcting their behavior (reducing the response rate compared to baseline) as time progressed.

In order to evaluate at what point of the devaluation (DV) test the animals actually displayed habitual behavior, we compared the cumulative devaluation index of the groups for each minute of the test against the hypothetical devaluation index of 0, indicating full habitual behavior, via one-sample t-test. During the Early time point, the control group showed goal-directed behavior throughout the entire test as the devaluation index was significantly different from 0 at all minutes. During the first minute of the test the CIE group displayed habitual behavior but goal-directed behavior throughout the rest of the DV test. During the Late time point, both control and CIE groups displayed habitual behavior during the first two minutes of the test but shift towards goal-directed behavior afterwards (**Figure 3D**, **Table 1**).

**Table 1.** One-sample t-test against 0 of the devaluation index for each cumulative minute of the extinction tests at early and late time points (Study 1, Experiment 2). Non-significant results indicate habitual behavior (highlighted in bold).

| Time point | Minutes (cumulative) | Controls |  |  | CIE |  |  |
| --- | --- | --- | --- | --- | --- | --- | --- |
| | | Mean ± SEM | t (df=11) | p-value | Mean ± SEM | t (df=8) | p-value |
| Early | 1 | 0.26 ± 0.09 | 2.854 | 0.016 | 0.04 ± 0.09 | 0,467 | **0.655** |
| | 2 | 0.37 ± 0.07 | 5.075 | 0.000 | 0.20 ± 0.07 | 2,688 | 0.028 |
| | 3 | 0.42 ± 0.07 | 5.842 | 0.000 | 0.18 ± 0.06 | 3,186 | 0.013 |
| | 4 | 0.45 ± 0.07 | 6.761 | 0.000 | 0.22 ± 0.06 | 3,411 | 0.009 |
| | 5 | 0.49 ± 0.06 | 8.329 | 0.000 | 0.26 ± 0.06 | 3,954 | 0.004 |
| Late | 1 | -0.06 ± 0.10 | -0.546 | 0.596 | -0.00 ± 0.10 | -0.031 | 0.976 |
| | 2 | 0.13 ± 0.09 | 1.495 | 0.163 | 0.06 ± 0.07 | 0.869 | 0.410 |
| | 3 | 0.24 ± 0.07 | 3.229 | 0.008 | 0.17 ± 0.07 | 2.672 | 0.028 |
| | 4 | 0.30 ± 0.06 | 5.427 | 0.000 | 0.23 ± 0.07 | 3.445 | 0.009 |
| | 5 | 0.35 ± 0.06 | 6.337 | 0.000 | 0.27 ± 0.07 | 3.766 | 0.005 |

### Study 2: Role of pDMS in spatial navigation and reward learning

We next used a DREADD-based approach to inhibit the pDMS in naïve Wistar rats and evaluated the effects on goal-directed and habitual behavior in both the spatial navigation and reward learning tasks (Experiment 3). We first trained the animals on the spatial navigation task, starting three weeks after the end of the last surgery in order to allow full expression of the transgenes, we then trained the same animals on the reward learning task starting one week after the end of the spatial navigation task (**Figure 1**). The quality of the injection of the virus was confirmed by immunohistochemistry for mCherry (expressing hM4Di, **Figure 4 A-B**) and GFP control virus within the pDMS region (**Figure 4 A**). Rats with misplaced virus injections were excluded from the entire analysis.

**Figure 4.**
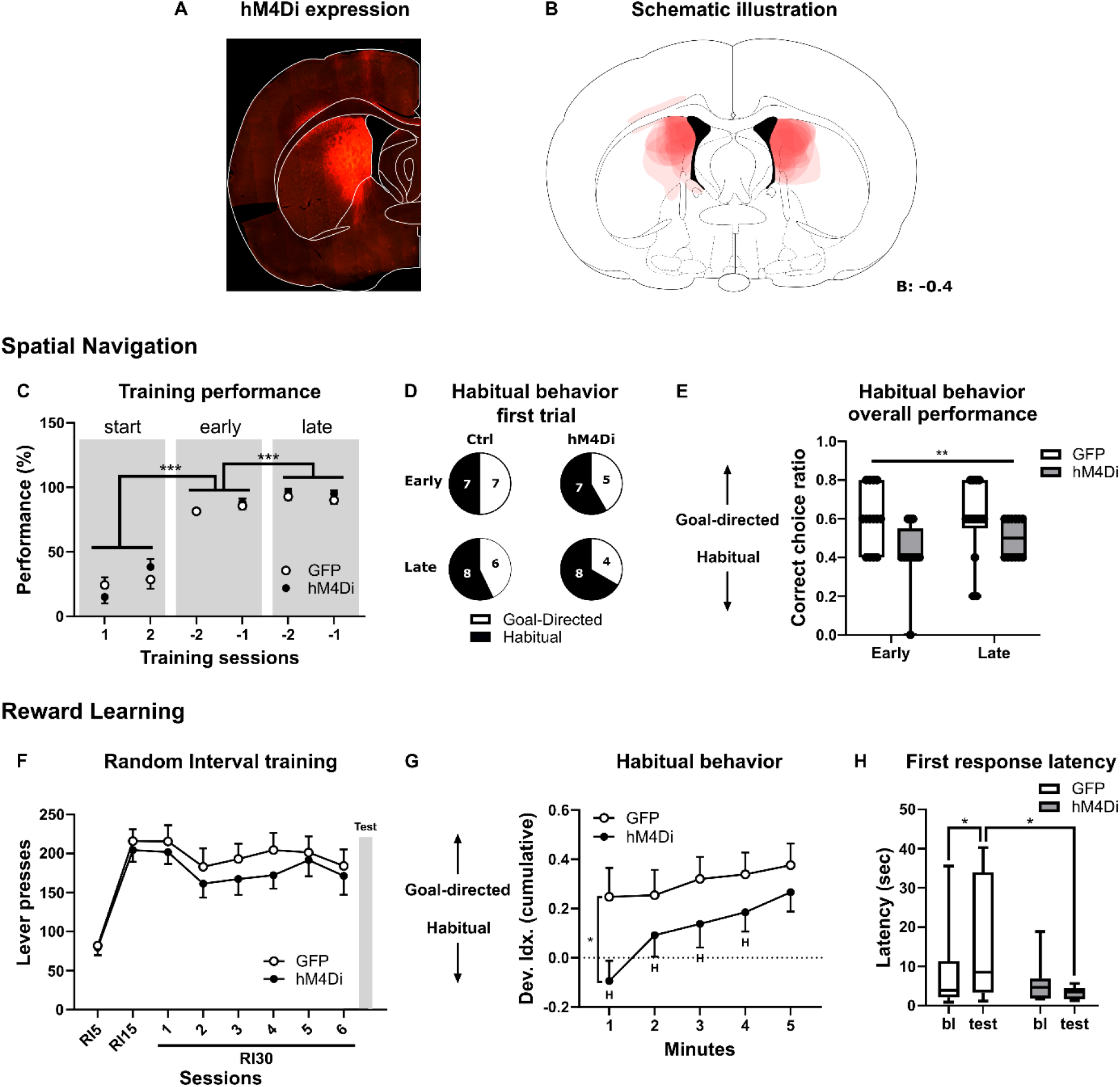
Inactivation of DMS led towards habitual behavior in naïve rats (Experiment 3). **A.** Coronal brain section showing the expression of hM4Di-mCherry within the pDMS. **B.** Schematic illustration showing quality of bilaterally injected hM4Di-mCherry virus at Bregma level: −0.4 mm. **C.** Training performance in percentage during the first two sessions, the last two sessions before the early probe session, and the last two session before the last probe session. **D.** Number of habitual choices made during the first trial for each probe session. **E.** Overall performance when considering all five trials for each probe session. **F.** Lever presses during random interval training. **G.** Devaluation index (Dev. Idx) over the course of the extinction test. A dev. Idx. of 0 indicates full habitual behavior, a Dev. Idx of 1 indicates full goal-directed behavior. Data points that were non-significantly different from 0 based on one-sample t-test are highlighted with “H”. Data in graphs **C**, **F**, and **G** are shown as mean ± SEM. Data in graph **D** are shown as pie chart. Data in graphs **E** and **H** are shown as box and whisker plot (median, first and third quartile range). Data from graph D were analyzed via χ^2^ test. Data from remaining graphs were analyzed via Repeated Measure Mixed Model. *p < 0.05, **p < 0.01, ***p < 0.001. DREADD- mediated inactivation of DMS was achieved by ip injection of CNO (Clozapine-N-oxide, 10mg/kg) 30 minutes prior each test. For Details, see Material & Methods.

### Spatial Navigation (T-Maze) task

As expected, we found no between groups difference (F_1,83_ = .5, p = .495) in performance during the training, but both groups progressively improved (F_2,89_ = 205, p = .000) going from the beginning of the training to prior the first probe session (p = .000) and from here to prior the second probe session (p = .000) (**Figure 4C**).

When analyzing the behavioral approach (habitual vs goal-directed) during the two probe sessions we did not find any difference between groups based on the first trial of each session (**Figure 4D**) as the hMD4i animals only made less than 20% more habitual turns during both time points, compared to GFP controls (early: χ^2^ = .2, p = .671; late: χ^2^ = .2, p = .619). However, when analyzing the overall performance considering all five trials of each probe session (**Figure 4E**) we found that hM4Di-treated rats displayed an overall increased number of habitual turns for both timepoints compared to GFP-expressing rats, as indicated by the presence of a statistically significant effect of Group (F_1,52_ = 9.9, p = .003) but lack of a Timepoint effect (F_1,52_ = .6, p = .424) and interaction (F_1,52_ = 1.3, p = .260) effect between the two factors. However, the post-hoc test indicated a significant difference between the groups at the Early (p = .004) but not at the Late (p = .163) time point.

### Reward learning (operant conditioning) task

After the end of the spatial navigation task, both groups were trained to self-administer 0.2% saccharin using the same FR1 pre-training used for Experiment 2 except that in this case the animals could not earn more than 50 rewards per session. This was done because we did not test the motivation of the animals in a progressive ratio test, because we had no reason to assume a change in the rewarding properties by the experimental procedure, and in order to reduce the risk for overtraining of the lever-pressing behavior. Once the random interval training started, all conditions were identical to the ones used in Experiment 2 but the animals were only tested once, after 6 RI-30 sessions, when goal-directed behavior would be expected. During RI training (**Figure 4F**) we found no difference in lever presses between the hM4Di and GFP groups (F_1,26_ = .7, p = .417).

For assessing habitual behavior, rats were treated with CNO (10 mg/Kg, ip.) 30 minutes before the test. When analyzing the behavior of the animals during the extinction test (**Figure 4G**), performed after reward satiety devaluation, we found a significant Group × Bin interaction (F_4,76_ = 5.7, p = .000) with the post-hoc test indicating that hM4Di-expressing rats had a significantly lower devaluation index, compared to controls, specifically during the first minute of the test (p = .011). Also in this case both groups corrected their behavior as the test progressed, as indicated by the significantly higher devaluation index during the first minute of the test and at the end of the test (Ctrl: p = .020; hM4Di: p = .000). Based on the one-sample t-test analysis of the devaluation index against 0, the hM4Di-expressing animals displayed habitual behavior during the first 4 minutes of the test, while control animals consistently displayed goal-directed behavior (**Figure 4G**, **Table 2**).

**Table 2.** One-sample t-test against 0 of the devaluation index for each cumulative minute of the DV test at the Early time point (Study 2, Experiment 3). Non-significant results indicate habitual behavior (highlighted in bold).

| Minutes (cumulative) | Ctrl |  |  | hM4Di |  |  |
| --- | --- | --- | --- | --- | --- | --- |
| | Mean ± SEM | t (df=11) | p-value | Mean ± SEM | t (df=7) | p-value |
| 1 | 0.26 ± 0.11 | 2.252 | 0.046 | -0.09 ± 0.08 | -1.149 | **0.294** |
| 2 | 0.25 ± 0.10 | 2.481 | 0.031 | 0.09 ± 0.09 | 1.047 | **0.335** |
| 3 | 0.32 ± 0.09 | 3.584 | 0.004 | 0.14 ± 0.10 | 1.438 | **0.201** |
| 4 | 0.34 ± 0.09 | 3.810 | 0.003 | 0.19 ± 0.08 | 2.340 | 0.058 |
| 5 | 0.38 ± 0.09 | 4.291 | 0.001 | 0.27 ± 0.08 | 3.379 | 0.015 |

We also analyzed the time required to make the first lever press response (first response latency, FRL) after the initiation of the extinction test and we compared it to the baseline FRL (**Figure 4H**), measured as the FRL averaged between the last two training sessions prior to the test. We found a significant Group × Time (baseline vs test) interaction (F_1,19_ = 5.1, p = .036) with the post-hoc test revealing that the control group took a longer time to make the first response during the test as compared to their baseline (p = .015), while the hM4Di animals did not (p = .438). Moreover, hM4Di-treated rats had a significantly lower FRL during the test compared to the response latency of the control animals during the test (p = .016).

## Discussion

In this study we provide evidence for a generalized habitization in alcohol dependent rats. We investigated habitual responses towards a sweet palatable reward in both tasks (spatial navigation and reward conditioning) and estimated to which extent such habitual responses are changed in abstinent rats. We found that most dependent rats displayed increased habitual responses at an earlier time point of learning, while non-dependent rats still showed a more goal-directed response. Chemogenetic inactivation of the pDMS in alcohol-naïve animals increases habitual tendencies and suggests alcohol-induced damage to this region in post-dependent rats. In conclusion, we confirm the critical role of DMS in maintaining goal-directed behavior, which seems to be to some extend compromised in AUD.

### Assessment of habitual behavior in the spatial navigation task

In study 1 we used post-dependent (CIE) and non-dependent rats for assessment of habitual behavior using a T-Maze navigation task. Both dependent and non-dependent controls displayed goal-directed responses at early time point of testing. Dependent rats switched to pronounced habitual responses after overtraining, while controls remained goal-directed (**Figure 2B-C**). The results imply that dependent rats developed habitual responses in a shorter time interval than controls, and overtraining was not effective in controls. A shift to habitual behavior in controls can be probably achieved after longer time of overtraining.

Navigation is an adaptive task in which stimulus-response associations or habits are incrementally acquired. Initially, a strategy based on a cognitive map (spatial map or ‘model-based’) are applied, while a more stereotyped response strategy (egocentric or ‘model-free’) can dominate after repeated training. Lesions of the dorsal striatum disrupt the latter strategy (Lee et al., 2008). AUD results in long-term damage of cortical regions, especially of the PFC but also hippocampus (Zahr et al., 2011; Meinhardt et al., 2013; Heilig et al., 2017; Meinhardt et al., 2021), suggesting that human subjects may preferentially adopt a striatal-based response strategy in e.g. 4-on-8 virtual maze or two step sequential Markov decision task.

The testing approach that we used in the spatial navigation task has one limitation: the five consecutive test trials during the early time point could influence the performance of the animals during the test performed at the late time point as the animals had the possibility to experience the test condition (inverted maze) multiple times, which would explain the increased overall performance in the control group from the early to the late time point.

### Assessment of habitual behavior in the operant conditioning task

Naïve rats were trained to self-administer 0.2% saccharin in operant chambers. Based on their motivation to self-administer saccharin, rats were assigned into alcohol dependent and control groups. After introduction of dependence by CIE, rats were trained for habitual behavior under a random interval (RI)-30 schedules that facilitated habitual behavior.

At the early and late time points rats were assessed for habitual behavior using a devaluation test procedure followed by an extinction test. At the early time point dependent rats displayed more habitual behavior (indicated by lower devaluation index) compared to controls. At the late time point both dependent and control rats showed a reduced devaluation index, indicating increased habitual behavior compared to the early time point (**Figure 3D**). Important to note, the dependent rats showed a higher motivation in the PR test for both saccharin and 10% alcohol during and at the end of the experiment, respectively, thus confirming the dependence-like phenotype (**Figure 3B**). An association between AUD and a sweet-liking phenotype has been in fact described in humans (Kampov-Polevoy et al., 2003; Pepino and Mennella, 2005, 2007; Bouhlal et al., 2018) and in high ethanol preferring rodents (Leventhal et al., 1995; Pecina and Berridge, 2005).

In the reward learning task, habitual behavior is typically tested in rodents after reward devaluation with an extinction test performed following a training under a random interval (RI) or random ratio (RR) schedule for a given number of sessions. Reward devaluation procedures, which can be accomplished via either specific satiety or conditioned-taste aversion, are typically used to dissociate habitual from goal-directed behavior (Dickinson, 1985; Barker and Taylor, 2014; McKim et al., 2016; Robbins and Costa, 2017). In this study the response to devaluation was assessed by pre-feeding the rats with saccharin solution prior to a 5-min extinction test. This test is usually short and it is performed in extinction conditions in order to avoid the animals experiencing the reward while in the devalued state and reduce extinction learning, this is particularly important if the animals are tested multiple times, as in our case. In this context, habitual behavior is defined as insensitivity to reward devaluation, meaning that animals that are displaying habitual behavior will press at a similar rate while in the devalued state (during the test) and when non-devalued (control condition/group) (Corbit et al., 2012). The length of the extinction test however is often different between studies that use this approach, lasting anywhere from 3 minutes to 30 minutes (Corbit et al., 2012; Mangieri et al., 2012; Clemens et al., 2014; Furlong et al., 2017; Iguchi et al., 2017). It is not clear if animals that show insensitivity to devaluation in a 3 minutes test would still show it in a 5 minutes test, or if animals found to be sensitive to the devaluation procedure in a 5 minutes test would also show sensitivity in a shorter test. Therefore, we have analyzed the data obtained from the devaluation-extinction test by calculating the devaluation index (Renteria et al., 2018; Towner and Spear, 2021) in cumulative form. The cumulative devaluation index is an estimate of how the behavior of the animals changes throughout the test.

In our experiment, following reward devaluation at the early time point, both dependent and control rats exhibited reductions in lever pressing, suggestive of a goal-directed response pattern. However, using the cumulative devaluation index we could show that dependent rats not only displayed overall increased habitual tendencies compared to controls (overall reduced devaluation index), but also that during the first minute of the test, dependent rats exhibited full habitual behavior, while control animals consistently exhibited goal-directed behavior throughout the test (**Figure 3D**). At the late time point, both dependent and control rats showed full habitual behavior during the first 2 minutes of the test while no longer differing in overall devaluation index, indicative of an effective overtraining.

A shift from goal-directed to habitual behavior is in line with the growing literature on dependence induced behavioral consequences, as there is evidence showing that chronic alcohol exposure can lead to increased habits also for taking behavior of a natural reward. Corbit and colleagues (2012) showed for example that non-contingent alcohol drinking performed during the course of operant training for a natural reward can induce insensitivity to outcome devaluation, while Towner and Spear (2021) showed that chronic intermittent ethanol intake, performed via oral gavage in late adolescent rats, also induces outcome insensitivity, indicating habitual behavior. Similarly, chronic intermittent alcohol vapor exposure has been shown to induce habitual behavior for a food reward in mice (Renteria et al., 2018). Interestingly, a recent study reported evidence of increased habitual behavior for taking of a non-alcohol reward also after a single acute dose of ethanol, however in alcohol-preferring mice (Houck and Grahame, 2018).

This body of literature adds up to the evidence that responding for alcohol transitions towards habitual behavior faster compared to responding for a natural reward (Dickinson et al., 2002; Corbit et al., 2012; Mangieri et al., 2012). Moreover, previous research demonstrated that food habits form more quickly in females than in males, but the opposite is true for alcohol habit formation (Quinn et al., 2007; Barker et al., 2010).

### Role of pDMS in spatial navigation and reward learning

Here we used a DREADD based approach to inhibit the posterior part of the dorsomedial striatum (pDMS) in naïve Wistar rats according to Yu et al., (2021). Inactivating DREADDs (hM4Di-mCherry) and GFP-expressing control viruses were bilaterally injected into the pDMS and effects on habitual behavior were assessed using both spatial navigation and operant tasks. We used a within subject design, with the same animals being tested for both behavior paradigms. In the T-Maze navigation task the DREADDs-mediated inactivation of the pDMS did induce an overall increased number of habitual choices at both time points, mostly pronounced during the early time point (**Figure 4E**). In the operant paradigm we could further demonstrate, that DREADD-treated rats displayed habitual behavior during the first 4 minutes of the devaluation test after short training, when goal-directed behavior would be expected, while the GFP-virus expressing control rats consistently displayed goal-directed behavior throughout the entire session. Thus, inactivation of pDMS by DREADDs led to habitual responses in both behavior paradigms, confirming the critical involvement of pDMS to maintain goal-directed behavior. Other studies found similar results after disengagement of the DMS and the formation of habitual behavior (Bassett et al., 2015; Kupferschmidt et al., 2017; Yu et al., 2021).

During the early goal-directed actions the DMS is mainly involved, while during the late habitual response, the DMS becomes weakened (Kupferschmidt et al., 2017) and the DLS instead becomes strengthened (Thorn et al., 2010). Thus, DMS and DLS seem to compete for control in the acquisition of habitual action sequences (Turner et al., 2022). Taken together, our findings support an opponent relationship of DMS and DLS in which loss of DMS function enhances the formation of habitual actions.

In general, instrumental responding (using an operant box) and spatial navigation (using a T-Maze) share important features. Firstly, two stages can be identified in the formation of a well-established action sequence: an early acquisition phase, characterized by a steep learning curve, and a phase of slow gradual consolidation, in which the behavior is optimized and becomes less susceptible to external influences. Secondly, the transition from early acquisition to later consolidation involves a shift from ventral to dorsal striatal regions. Control of motivated behaviors is initiated in the ventral striatum. Clinical studies suggest that during the transition from reward or goal-directed to habit-driven behavior, control is shifted towards dorsal parts of the striatum (Vollstadt-Klein et al., 2010). The most direct evidence for this process comes from a series of studies, which directly showed ventral striatum activation to alcohol challenge in light social drinkers, attenuation of this signal in heavy social drinkers (some of whom met criteria for diagnosis), and lack of this response in hospitalized treatment-seeking patients (Gilman et al., 2008; Gilman et al., 2012; Spagnolo et al., 2014).

There are human fMRI studies showing a reduced sensitivity to devaluation after extensive instrumental training (Tricomi et al., 2009; Sjoerds et al., 2013), but other studies did not find evidence for habitual responses in human subjects after prolonged training (de Wit et al., 2009; Hogarth et al., 2019; Hogarth and Field, 2020; Luijten et al., 2020). Acute treatment of alcohol had been shown to impair goal-directed action (Hogarth et al., 2012). First evidence for a shift from goal-directed to habitual behavior in AUDs came from Sjoerds et al. (2013) using an instrumental learning task. Later on, another study compared AUD patients with healthy controls using the two-stage Markov decision task and found a reduction of goal-directed performance in AUDs, but no difference in habitual responding (Sebold et al., 2014). This finding may highlight rather the possibility of general cognitive impairments in AUDs that impact of habitual action control. Other studies using the same task compared a variety of compulsive disorders including abstinent AUD patients (Voon et al., 2015). Similarly to Sebold et al. (2014), they did not observe whole-sample differences between AUD patients and healthy controls. However, the study showed an association of early abstinence with greater habitual action control which shifts towards more goal-directed control with longer abstinence. Another study using a web-based version of the two-stage Markov decision task with 1413 participants showed a significant decrease in goal-directed control in participants with self-reported AUD (using AUDIT) which were attributed to impairment in goal-directed control to a broader range of trans-diagnostic factors, which they grouped as compulsive behavior and intrusive thought (Gillan et al., 2016). A recent longitudinally designed study with a sample size of 188 young social drinkers found that the amount of regular alcohol consumption had no significant impact on action control and habits (Nebe et al., 2018). All these contradictory findings contrast the results on habitual overreliance in AUD patients (Tricomi et al., 2009; Sjoerds et al., 2013), which might be due to the different methodologies, sample features and relatively smaller sample sizes in the majority of studies.

In our animals, habitual behavior was not detected in rats with low withdrawal scores (data not shown) or least AUD-like behavior in terms of extensive homecage drinking (Smeets et al., 2022). Thus, a behavior heterogeneity of the AUD-like phenotype, such as e.g. amount of alcohol consumption/self-administration, number of lever presses/motivation, withdrawal severity, sweet-liking phenotype (Bouhlal et al., 2018), seem to be important output factors for assessment of habitual behavior in rodents. In addition, we observed variability and low effects sizes of the observed effects. A large individual variability in compulsivity has been observed in animals self-administrating drugs such as cocaine and alcohol (Giuliano et al., 2019; Siciliano et al., 2019).

Further potential limitations of the study were that we did not consider habitual behavior for alcohol reward and other aspects of the AUD-like phenotype such as e.g. loss of control for aversive-/footshock-resistant responding for alcohol- or saccharin reward. Finally, it is important to identify molecular mechanism of habitual behavior for natural rewards to better understand how this behavior developed in comparison with alcohol rewarding effects.

In summary, we conclude that dependence induced a generalized habitual response in rats for both spatial navigation and reward conditioning tasks. Inactivation of pDMS led to habitual responses, similar as seen in our dependent rats. Thus, pDMS is critical to maintain goal-directed behavior; modulation of its activity will lead to changes in habitual responses.

Based on our findings and a growing body of evidence we conclude a more minor role of habit formation in regard to AUD-like behavior. However, our data provide potential implications for understanding how dorsal striatal dysfunction contributes to AUD, and thus may provide new therapeutic strategies in support for regaining control over ‘bad habits’.

## Acknowledgments

Financial support for this work was provided by the Bundesministerium für Bildung und Forschung (BMBF) funded SysMedSUDs consortium (FKZ: 01ZX1909A), the Deutsche Forschungsgemeinschaft (DFG, German Research Foundation—Project-ID: ME 5279/3-1 and Project-ID 402170461—TRR 265 (Heinz et al., 2020)) as well as the Hetzler Foundation for Addiction Research.

## Author Contributions

ACH, RS, and WHS were responsible for study design and procured study funding. FG, AH, MC, MMF, and MM performed the behavioral experiments. FG analyzed data. FG, RS, WHS, and ACH wrote the manuscript.

